# Clustering of Type 2 Diabetes Genetic Loci by Multi-Trait Associations Identifies Disease Mechanisms and Subtypes

**DOI:** 10.1101/319509

**Authors:** Miriam S. Udler, Jaegil Kim, Marcin von Grotthuss, Sílvia Bonàs-Guarch, Josep M Mercader, Joanne B. Cole, Joshua Chiou, Christopher D Anderson, Michael Boehnke, Markku Laakso, Gil Atzmon, Benjamin Glaser, Kyle Gaulton, Jason Flannick, Gad Getz, Jose C. Florez

**Author notes:** **Corresponding author:** Dr. Jose C. Florez, Center for Genomic Medicine Massachusetts General Hospital, Simches Research Building 185 Cambridge Street, CPZN-5 Boston, MA 02114.

## Abstract

**Background:** Type 2 diabetes (T2D) is a heterogeneous disease for which 1) disease-causing pathways are incompletely understood and 2) sub-classification may improve patient management. Unlike other biomarkers, germline genetic markers do not change with disease progression or treatment. In this paper we test whether a germline genetic approach informed by physiology can be used to deconstruct T2D heterogeneity. First, we aimed to categorize genetic loci into groups representing likely disease mechanistic pathways. Second, we asked whether the novel clusters of genetic loci we identified have any broad clinical consequence, as assessed in four independent cohorts of individuals with T2D.

**Methods and Findings:** In an effort to identify mechanistic pathways driven by established T2D genetic loci, we applied Bayesian nonnegative matrix factorization clustering to genome-wide association results for 94 independent T2D genetic loci and 47 diabetes-related traits. We identified five robust clusters of T2D loci and traits, each with distinct tissue-specific enhancer enrichment based on analysis of epigenomic data from 28 cell types. Two clusters contained variant-trait associations indicative of reduced beta-cell function, differing from each other by high vs. low proinsulin levels. The three other clusters displayed features of insulin resistance: obesity-mediated (high BMI, waist circumference), “lipodystrophy-like” fat distribution (low BMI, adiponectin, HDL-cholesterol, and high triglycerides), and disrupted liver lipid metabolism (low triglycerides). Increased cluster GRS’s were associated with distinct clinical outcomes, including increased blood pressure, coronary artery disease, and stroke risk. We evaluated the potential for clinical impact of these clusters in four studies containing participants with T2D (METSIM, N=487; Ashkenazi, N=509; Partners Biobank, N=2,065; UK Biobank N=14,813). Individuals with T2D in the top genetic risk score decile for each cluster reproducibly exhibited the predicted cluster-associated phenotypes, with ~30% of all participants assigned to just one cluster top decile.

**Conclusion:** Our approach identifies salient T2D genetically anchored and physiologically informed pathways, and supports use of genetics to deconstruct T2D heterogeneity. Classification of patients by these genetic pathways may offer a step toward genetically informed T2D patient management.

## Introduction

Type 2 diabetes (T2D) is a complex disease affecting the world’s population at epidemic rates and whose pathophysiology remains incompletely understood. Approximately 30.3 million (9.4%) of people in the United States have diabetes, with T2D thought to account for 90–95% of all diagnoses (1, 2). Despite recognized heterogeneity in patient phenotypes and responses to treatment, T2D management strategies remain largely impersonalized.

In an attempt to deconstruct the heterogeneity of T2D, recent studies have performed cluster analysis of individuals using serum biomarkers and clinical data to identify T2D subgroups (3, 4). These studies offer exciting directions for future research, but are also limited by the nature of the variables included in analyses. For example Ahlqvist *et al*. (4) clustered individuals using six variables measured shortly after diabetes diagnosis, including hemoglobin A1c, glutamate decarboxylase antibodies, and body mass index (BMI); notably, these variables change with disease progression and treatment, and thus application of this clustering approach to clinical practice is of uncertain utility when patients are evaluated at a different time in the disease course or after treatment has been initiated. Additionally, it is not clear whether clinical biomarkers used in clusters analyses to date are causal, consequential, or coincidental in the disease process.

In contrast to other serum biomarkers, germline genetic variants associated with T2D are more likely to point to T2D causal mechanisms and remain constant regardless of developmental stage, disease state, or treatment. Over the past decade, genome-wide association studies (GWAS) and other large-scale genomic studies have identified over 100 loci associated with T2D, causing modest increases in disease risk (odds ratios generally < 1.2) (5–9). These genetic loci offer insight into biological pathways causing T2D, but for most of these loci the causal variant(s) and the mechanism by which the locus causes T2D remain unknown, limiting opportunities for clinical translation.

As GWAS have been conducted across multiple traits, there exists an opportunity to leverage multi-variant-trait association patterns to elucidate likely shared disease mechanisms, based on the assumption that genetic variants that act along a shared pathway will have similar directional impact on various observed traits. For example, amongst genetic variants impacting insulin resistance, Yaghootkar *et al*. identified a set of 11 variants associated with a particular directional pattern of traits in GWAS. This set of 11 insulin-resistance-increasing alleles was felt to represent a “lipodystrophy-like” fat distribution subgroup of insulin resistance variants reminiscent of monogenic lipodystrophy, since they were associated with increased fasting insulin and triglyceride levels, but decreased HDL cholesterol, adiponectin, and BMI (10).

As with insulin resistance, T2D-associated genetic variants have been assessed using a similar multi-variant-trait clustering approach, however, the resultant clusters have had limited clinical interpretability to date. Three efforts to perform clustering of T2D loci have been published by Dimas *et al*. (11) focusing on glycemic traits, and recently by Scott *et al*. (6) and Mahajan *et al*. (9), both including BMI and lipid traits in addition to glycemic traits. In these analyses, unsupervised hierarchical clustering was performed on T2D variants using their associations with respective traits. While these approaches generated some biologically suggestive clusters of genetic loci, determining the number and boundaries of clusters using unsupervised hierarchical clustering remains rather subjective. Additionally, in these analyses, many variants could not be clustered (more than half of all loci included in (6, 11)), including loci with known mechanism, tissue specificity, and physiological impact (e.g. those in *HNF1A* and *TM6SF2*). The unsupervised hierarchical clustering model applied in these previous efforts requires that a variant be included in only one pathway, so-called “hard-clustering,” and were limited by the GWAS datasets available for diabetes-related traits. We were interested in investigating a more flexible model that would allow a variant to impact more than one biological pathway and hypothesized that this might improve cluster interpretability, using the most up-to-date GWAS datasets available for metabolic traits.

In this paper we test whether a germline genetic approach can be used to deconstruct T2D heterogeneity. First, we ask whether genetic variants can be categorized into groups representing likely disease mechanistic pathways. We apply the novel clustering method Bayesian non-negative matrix factorization (bNMF), to enable a “soft-clustering”, whereby a variant can be associated to more than one cluster, which has been used previously in cancer genomics (12–15). To confirm that these groups of variants represent distinct entities with predicted biological relevance, we assess for tissue-specificity for enhancer or promoter enrichment. Second, we ask whether the novel clusters of genetic loci we identified are of any clinical consequence, which we assess in four independent cohorts of individuals with T2D.

## Methods

### Variant and trait selection

To obtain a comprehensive set of genetic variants associated with T2D, we started with the set of 88 variants reaching genome-wide significance aggregated by Mohlke and Boehnke (5), and then added 37 additional loci that were reported in subsequent T2D large-scale genetic studies (6, 7, 9). At some loci, there have been reports of multiple distinct signals (6, 8); we included 9 additional variants representing distinct signals at 6 loci (*ANKRD55, DGKB, CDKN2A, KCNQ1, CCND2*, and *HNF4A*). Of the 125 T2D loci considered, we selected a subset of 94 representative variants based on the condition that either the variant or a proxy of the variant had an association with T2D in the DIAGRAM version 3 (DIAGRAMv3) Stage 1 meta-analysis (16) with *P*<0.05 **(Table S1)**. Proxies of variants were required to either be in linkage disequilibrium (r^2^ > 0.6) with an original T2D variant published at genome-wide significance, be the lead SNP at the T2D locus in DIAGRAMv3, or reach genome-wide significance in DIAGRAMv3. Since DIAGRAMv3 contains study populations of mostly European ancestry, focusing on variants with at least nominal significance in this dataset helped ensure that these variants would also be associated with other traits in published GWAS, as most of the GWAS populations were also of predominantly European ancestry. Additionally, variants (other than those representing distinct signals at a locus) were conservatively excluded if they fell within 500 kb of other another variant on the list. Given that C/G and A/T alleles are ambiguous and can lead to errors in aligning alleles across GWAS, as the analyses progressed, we opted to avoid inclusion of ambiguous alleles, choosing proxies instead.

For the 94 T2D-associated variants, the T2D-risk increasing alleles were identified and all future analyses used the aligned T2D risk-increasing alleles. Summary association statistics for additional traits whose GWAS meta-analyses were publicly available were aggregated for each variant **(Table S2)**. These traits included glycemic traits available through the Meta-Analyses of Glucose and Insulin-related traits Consortium (MAGIC) (fasting insulin, fasting glucose, fasting insulin adjusted for BMI, 2-hour glucose after oral glucose tolerance test [OGTT] adjusted for BMI, glycated haemoglobin [HbA1c], homeostatic model assessments of beta-cell function [HOMA-B] and insulin resistance [HOMA-IR], incremental insulin response at 30 minutes on OGTT, insulin secretion at 30 minutes on OGTT, fasting proinsulin adjusted for fasting insulin, corrected insulin response [CIR], disposition index, and insulin sensitivity index [ISI]) (17–23). Anthroprometric traits were publicly available through the Genetic Investigation of ANthrometric Traits (GIANT) consortium (BMI, height, waist circumference [WC] with and without adjustment for BMI, waist-hip ratio [WHR] with and without adjustment for BMI) (24–26). Additional birth weight and length GWAS summary statistics were obtained from the Early Growth Genetics Consortium (EGG) consortium (27, 28). GWAS results for visceral and subcutaneous adipose tissue were available from VATGen consortium (29), as well as results for percent body fat (30) and heart rate (31). Finally serum laboratory values were available for the following traits: lipid levels (HDL cholesterol, LDL cholesterol, total cholesterol, triglycerides) (32), leptin with and without BMI adjustment (33), adiponectin adjusted for BMI (34), urate (35), N3-fatty acids (36), N6-fatty acids (37), plasma phospholipid fatty acids in the de novo lipogenesis pathway (38), and very long-chain saturated fatty acids (39). To reduce noise, traits were only included if there was at least one variant was associated with the trait at a Bonferroni-corrected threshold of significance *P*<5×10^−4^ (0.05/94).

In addition to the above traits used in the clustering process, single-variant association results with ten clinical outcomes were also aggregated **(Table S2)**: ischemic stroke and its subtypes (large vessel disease, small vessel disease, and cardioembolic) (40), coronary artery disease (41), renal function as defined by estimated glomerular filtration rate (eGFR) and urine albumin-creatinine ratio (UACR), and chronic kidney disease (CKD) (42), and systolic and diastolic blood pressure (43).

### Bayesian Non-negative Matrix Factorization clustering

Z-scores for variant trait associations from GWAS were generated by dividing beta by the standard error, using the summary statistic results. To address the marked differences in sample size across studies and allow for a more uniform comparison of phenotypes across studies, the z-scores were scaled by the square root of study size, as estimated by mean sample size across all SNPs, forming the variant-trait association matrix ***Z*** (94 by 47).

To enable an inference for latent overlapping modules or clusters embedded in variant-trait associations we modified the existing bNMF algorithm to explicitly account for both positive and negative associations. Each column of ***Z*** split into two separate entities where one contained all positive z-scores, the other has all negative z-scores multiplied by −1, and setting zero otherwise, leading to the association matrix ***X*** (94 by 94) comprised of doubled traits with positive and negative associations to variants. Then bNMF factorizes ***X*** into two matrices, ***W*** (94 by *K*) and ***H***^T^ (94 by *K*) with an optimal rank *K*, as ***X*** ~ ***WH***, corresponding the association matrix of variants and traits to the determined clusters, respectively. This mathematical framework enables bNMF to tackle both positive and negative associations with no loss of information, while keeping its non-negativity constraint. Determining the proper model order *K* is a key aspect in balancing data fidelity and complexity. Conventional NMF requires the model order as an input or it may be determined post data-processing, but bNMF is designed to suggest an optimal *K* best explaining ***X*** at the balance between an error measure, ||X-WH||^2^, and a penalty for model complexity derived from a non-negative half-normal prior for ***W*** and ***H*** (12–14). The defining features of each cluster are determined by the most highly associated traits, which is a natural output of the bNMF approach. bNMF algorithm was performed in R Studio for 1000 iterations with different initial conditions and the most probable solutions for *K* was selected for downstream analysis. The results of clustering provides cluster-specific weights for each variant (***W***) and trait (***H***).

### Trait and outcome associations with each cluster

Associations of the GRS for each cluster with each GWAS trait or outcome was performed using inverse-variance weighted fixed effects meta-analysis using summary statistics from GWAS, as has been done previously (10). The associations with GRS’s and traits were performed to confirm clustering results; outcomes were not included in the clustering process. For these analyses, the top set of strongest weighted variants for each cluster were included in the model using a cut off weighting of 0.75, which was determined by two independent approaches involving modeling of cluster weights. First the top-weighted trait in each cluster was assessed with variants in the corresponding cluster in a step-wise approach; a meta-analysis of the variants with the top-weighted trait was performed, starting with all variants and removing sequential variants from lowest to highest weighted until the local minimum meta-analysis *P*-value was obtained. As a second approach, we assessed the distribution of the variant clustering weights for each variant across all clusters with the goal of identifying optimal cut-offs to define the beginning of the “long tail,” representing less informative variants for each cluster **(Figure S3)**. We plotted the delta of consecutive clustering weights sorted in descending order and reasoned that the long tail should start just after the last significant difference in consecutive weights **(Figure S4)**. Therefore, all clustering weight deltas were order in descending order, and the top 5% were considered to be significant deltas. The last significant delta is shown in **Figure S4** with an arrow, and it corresponds to clustering weight of 0.75.

Ten outcomes were assessed, which were independent from the traits included in the bNMF clustering. A conservative significance threshold was set at 1×10^−3^ using a Bonferroni correction for ten traits and 5 clusters (0.05/50).

### Functional annotation and enrichment analysis

We calculated Bayes Factors for 100% credible set variants at each locus from effect size estimates and standard errors using the approach of Wakefield (44). We then calculated a posterior probability for each variant by dividing the Bayes Factor by the sum of all Bayes Factors in the credible set. We obtained previously published 13-state ChromHMM (45) chromatin state calls for 28 cell-types excluding cancer cell lines (46). For each cell type, we extracted chromatin state annotations for enhancer (Active Enhancer 1, Active Enhancer 2, Weak Enhancer, Genic Enhancer) and promoter (Active Promoter) elements.

We assessed enrichment of annotations first within clusters and second across clusters. For within cluster analysis, we overlapped cell-type annotations with credible set variants for loci in each cluster. We then calculated a cell-type probability for the cluster as the sum of posterior probabilities of variants in cell-type enhancers or promoters divided by the number of loci in the cluster. For the across-cluster analysis, we overlapped cell-type annotations with credible set variants for all loci. For each cluster, we then calculated a cell type probability as the sum of posterior probabilities of all variants in cell type enhancers or promoters in the cluster divided by the total number of loci in the cluster.

We derived significance for cell type probabilities for each cluster using a permutation-based test. Within each cluster, we permuted locus and cell-type labels and then recalculated cell-type probabilities as above. For across-cluster analysis, we permuted cluster labels for each locus, and then recalculated cell type probabilities for the permuted clusters as above. For the 5 loci (*ADCY5*, *CDC123*, *HNF4A*, *HSD17B12*, *CCND2*) represented in multiple clusters, we ensured that each locus was only represented once per cluster. We then used the cell-type probabilities derived from 1M permutations as a background distribution and performed a one-tailed test to ascertain significance for each cell type.

## Study Populations

### Metabolic Syndrome in Men Study (METSIM)

From this cross-sectional study of Finnish men (47), we analyzed data from 487 individuals with T2D previously ascertained for genotyping as part of the T2D-GENES initiative (48). Genotyping was preformed using Illumina HumanExome-12v1_A Beadchip, and imputation was performed using the Haplotype Consortium Reference Panel (49) using the Michigan Imputation Server (50).

### Diabetes Genes in Founder Populations (Ashkenazi) study

We analyzed data from 509 individuals with T2D from this study previously ascertained for genotyping as part of the T2D-GENES initiative (48). Briefly, the study consists of individuals of Ashkenazi Jewish origin selected from two separate DNA collections: 1. One affected individual selected per family from a genome-wide, affected-sibling-pair linkage study (51). Families in which both parents were known to have diabetes were excluded. 2. Patients ascertained by the Israel Diabetes Research Group between 2002 and 2004 from 15 diabetes clinics throughout Israel. For this study, only T2D patients with age of diagnosis between 35 and 60 were selected (52). Genotyping was performed using Illumina Cardio-Metabo Chip and imputation was performed using the Haplotype Consortium Reference Panel (49) using the Michigan Imputation Server (50).

### The Partners Biobank

The Partners HealthCare Biobank maintains blood and DNA samples from more than 60,000 consented patients seen at Partners HealthCare hospitals, including Massachusetts General Hospital, Brigham and Women’s Hospital, McLean Hospital, and Spaulding Rehabilitation Hospital, all in the Boston area, Massachusetts (53). Patients are recruited in the context of clinical care appointments at more than 40 sites and clinics, and also electronically through the patient portal at Partners HealthCare. Biobank subjects provide consent for the use of their samples and data in broad-based research. The Partners Biobank works closely with the Partners Research Patient Data Registry (RPDR), the Partners’ enterprise scale data repository designed to foster investigator access to a wide variety of phenotypic data on more than 4 million Partners HealthCare patients. Approval for analysis of Biobank data was obtained by Partners IRB, study 2016P001018.

Type 2 diabetes status was defined based on “curated phenotypes” developed by the Biobank Portal team using both structured and unstructured electronic medical record (EMR) data and clinical, computational and statistical methods (54). Cases were selected by this algorithm to have type 2 diabetes with PPV of 90% and required to be at least age 35 in order to further minimize misclassification of T2D diagnosis. Additional phenotypic data was extracted using the most recent value within the past five years. Genomic data for 15,061 participants was generated with the Illumina Multi-Ethnic Genotyping Array. The genotyping data was harmonized and quality controlled with a 3-step protocol, including two stages of SNP removal and an intermediate stage of sample exclusion. The exclusion criteria for variants was: (i) missing call rate ≥0.05, (ii) significant deviation from Hardy-Weinberg equilibrium (*P*≤1×10^−6^ for controls and *P*≤1×10^−20^ for the entire cohort), (iii) significant differences in the proportion of missingness between cases and controls *P*≤1×10^−6^ and (iv) minor allele frequency <0.001. The exclusion criteria for samples was: i) gender discordance between the reported and genetically predicted sex, ii) subject relatedness (pairs with ≥0.125 from which we removed the individual with the highest proportion of missingness), iii) missing call rates per sample ≥0.02 and iv) population structure showing more than 4 standard deviations within the distribution of the study population according to the first four principal components. Phasing was performed with SHAPEIT2 (55) and then imputed with the Haplotype Consortium Reference Panel (49) using the Michigan Imputation Server (50).

### The UK Biobank

UK Biobank (UKBB) is a prospective cohort of ~500,000 recruited participants from the general population aged 40–69 years in 2006–2010 from across the UK, with genotype, phenotype and linked healthcare record data. Individuals in UKBB underwent genotyping with one of two closely related custom arrays (UK BiLEVE Axiom Array or UK Biobank Axiom Array) consisting of over 800,000 genetic markers scattered across the genome. Additional genotypes were imputed centrally using the Haplotype Reference Consortium (HRC) reference panel (49), 1000G phase 3 (56), and UK10K reference panel (57), as previously reported (58). All the SNPs used for computing the GRS were imputed only with the HRC reference panel. We restricted the analysis to a subset of unrelated individuals of white European ancestry, constructed centrally using a combination of self-reported ancestry and genetically confirmed ancestry by projecting UKBB genetic principal components on to 1000G phase 3 reference principal component space. We focused on individuals with T2D, based on a recently developed algorithm of information at the baseline visit that took into account a nurse interview self-reported diagnosis, diabetes medication use, and age at diagnosis (59). We expanded upon this definition to include touchscreen self-reported diagnosis, diagnosis and medication information provided at repeat visits, and removed individuals with reported “age of diabetes diagnosis” less than 40 to further reduce possible contamination with type 1 diabetes diagnoses. 14,813 individuals determined by the algorithm to have “probable” or “possible” T2D were included in our analyses.

### Individual-level analyses of individuals with T2D

Individual-level analyses were performed using data from METSIM, the Diabetes Genes in Founder Populations (Ashkenazi) study, the Partners Biobank, and UK Biobank. Analyses were restricted to individuals with T2D and Caucasian ancestry.

All SNPs were genotyped or imputed with high quality (Rsq values > 0.95). SNPs were included in genetic risks scores as allele dosages. GRS’s were generated for each cluster by multiplying a variant’s genotype dosage by its cluster weight. Only the top-weighted variants falling above the threshold, as defined above, were included in the GRS for each cluster. Logistic regression and linear regression performed in Stata v14.2 adjusting for sex, age, and principal components.

Results from the multiple cohorts were meta-analyzed using an inverse-variance weighted fixed effect model. Traits in subgroups of individuals with top decile cluster GRS’s were compared using the Kruskal-Wallis test for continues traits, except for percentage female sex, which was compared with the Chi-squared test.

### Results

### Clustering suggests five dominant pathways driving diabetes risk

Clustering of variant-trait associations was performed for 94 genetic variants and 47 traits derived from GWAS using bNMF clustering, with identification of five robust clusters present on 82.3% of iterations **(Figures S1, S2; Tables S1–4)**. In 17.3% of the iterations the data converged to four clusters, in which Cluster 2 was subsumed into Cluster 1. As bNMF clusters both variants and traits, the top-weighted traits can be used to help define the underlying mechanism in each cluster. The five clusters appeared to represent two mechanisms of beta-cell dysfunction and three mechanisms of insulin resistance (Figure 1a, Table 1, **Table S5**).

**Table 1.** Associations of cluster genetic risk scores and selected GWAS traits

|  | Beta Cell<br>N Loci = 30 |  | Proinsulin<br>N Loci = 7 |  | Obesity<br>N Loci = 5 |  | Lipodystrophy<br>N Loci = 20 |  | Liver/Lipid<br>N Loci = 5 |  |
| --- | --- | --- | --- | --- | --- | --- | --- | --- | --- | --- |
| Trait | beta | Pvalue | beta | Pvalue | beta | Pvalue | beta | Pvalue | beta | Pvalue |
| Adiponectin | -0.0005 | 0.55 | -0.0019 | 0.37 | **-0.0007** | 0.74 | **-0.0114** | **3.34E<sup>-23</sup>** | -0.0007 | 0.77 |
| BMI | **-0.0026** | **6.0x10<sup>-5</sup>** | **-0.0080** | **3.1x10<sup>-8</sup>** | **0.0396** | **9.7x10<sup>-157</sup>** | **-0.0079** | **1.81E<sup>-21</sup>** | 0.0001 | 0.94 |
| Bodyfat | -0.0016 | 0.11 | -0.0061 | 4.5x10<sup>-3</sup> | **0.0247** | **2.1x10<sup>-25</sup>** | **-0.0120** | **9.04E<sup>-22</sup>** | -0.0031 | 0.26 |
| CIR | **-0.0584** | **7.1x10<sup>-43</sup>** | -0.0234 | 0.014 | -0.0010 | 0.92 | 0.0087 | 0.10 | -0.0021 | 0.85 |
| DI | **-0.0543** | **6.6x10<sup>-37</sup>** | -0.0080 | 0.40 | -0.0086 | 0.40 | -0.0102 | 0.05 | -0.0115 | 0.30 |
| 2hrGlu | **0.0288** | **2.0x10<sup>-13</sup>** | 0.0204 | 0.02 | 0.0064 | 0.49 | **0.0292** | **2.26E<sup>-9</sup>** | -0.0257 | 0.01 |
| FI | **-0.0033** | **4.4x10<sup>-7</sup>** | -0.0054 | 2.2x10<sup>-4</sup> | **0.0087** | **6.1x10<sup>-8</sup>** | **0.0068** | **1.96E<sup>-16</sup>** | **0.0071** | **3.8x10<sup>-5</sup>** |
| FI adj BMI | **-0.0026** | **4.4x10<sup>-6</sup>** | -0.0040 | 1.3 x10<sup>-3</sup> | -0.0008 | 0.57 | **0.0082** | **3.01E<sup>-31</sup>** | **0.0082** | **2.1x10<sup>-8</sup>** |
| HDL | -0.0008 | 0.51 | -0.0031 | 0.27 | -0.0059 | 0.05 | **-0.0191** | **1.96E<sup>-33</sup>** | 0.0069 | 0.038 |
| Height | 0.0009 | 0.12 | -0.0058 | **3.3x10<sup>-5</sup>** | **-0.0033** | **1.9x10<sup>-5</sup>** | **0.0061** | **4.44E<sup>-05</sup>** | -0.0005 | 0.77 |
| HC | -0.0031 | **3.5x10<sup>-5</sup>** | **-0.0113** | **1.0x10<sup>-11</sup>** | 0.0345 | **9.1x10<sup>-79</sup>** | **-0.0116** | **9.69E<sup>-34</sup>** | -0.0007 | 0.73 |
| HOMA-B | **-0.0066** | **1.9x10<sup>-21</sup>** | **-0.0103** | **2.6x10<sup>-11</sup>** | 0.0066 | **8.0x10<sup>-5</sup>** | 0.0019 | 0.03 | 0.0019 | 0.30 |
| HOMA-IR | -0.0011 | 0.21 | -0.0041 | 0.03 | **0.0108** | **9.0x10<sup>-8</sup>** | **0.0066** | **2.2x10<sup>-10</sup>** | **0.0093** | **2.6x10<sup>-5</sup>** |
| Incr30 | **-0.0398** | **6.9x10<sup>-19</sup>** | -0.0239 | 0.02 | -0.0053 | 0.63 | 0.0198 | 4.8x10<sup>-4</sup> | 0.0102 | 0.38 |
| Ins30_bmi | **-0.0503** | **1.8x10<sup>-28</sup>** | -0.0310 | 1.8x10<sup>-3</sup> | 0.0027 | 0.81 | 0.0163 | 3.9x10<sup>-3</sup> | 0.0054 | 0.64 |
| ISI adj BMI | -0.0039 | 0.06 | -0.0020 | 0.67 | 0.0045 | 0.37 | **-0.0213** | **1.3x10<sup>-13</sup>** | -0.0086 | 0.12 |
| Leptin | 0.0009 | 0.50 | -0.0067 | 0.03 | **0.0197** | **1.0x10<sup>-9</sup>** | **-0.0245** | **8.9x10<sup>-21</sup>** | **0.0147** | **3.1x10<sup>-5</sup>** |
| Linoleic acid | 0.0093 | 0.29 | -0.0232 | 0.25 | 0.0027 | 0.90 | -0.0024 | 0.83 | **0.1330** | **1.31E-8** |
| Palmitoleic | 0.0002 | 0.74 | 0.0024 | 0.11 | 0.0034 | 0.03 | -0.0020 | 0.02 | **-0.0104** | **5.50E-9** |
| Proinsulin | **0.0097** | **1.2x10<sup>-10</sup>** | -0.0297 | **1.4x10<sup>-18</sup>** | 0.0047 | 0.18 | 0.0059 | 1.3x10<sup>-3</sup> | 0.0059 | 0.13 |
| Total Chol | 0.0023 | 0.06 | -0.0055 | 0.04 | -0.0023 | 0.45 | 0.0046 | 3.2x10<sup>-3</sup> | **-0.0182** | **3.1x10<sup>-8</sup>** |
| Triglycerides | 0.0022 | 0.07 | -0.0027 | 0.33 | 0.0066 | 0.03 | **0.0194** | **1.8x10<sup>-34</sup>** | **-0.0416** | **1.0x10<sup>-35</sup>** |
| Urate | -0.0007 | 0.51 | -0.0045 | 0.084 | **0.0165** | **1.4x10<sup>-9</sup>** | **0.0090** | **2.2x10<sup>-10</sup>** | **-0.0260** | **2.6x10<sup>-18</sup>** |
| WC | -0.0020 | 5.23x10<sup>-3</sup> | **-0.0096** | **1.5x10<sup>-9</sup>** | **0.0379** | **1.0x10<sup>-102</sup>** | **-0.0058** | **3.6x10<sup>-10</sup>** | -0.0005 | 0.80 |
| WC female | -0.0010 | 0.30 | -0.0073 | 3.9x10<sup>-4</sup> | **0.0376** | **1.4x10<sup>-60</sup>** | -0.0022 | 0.07 | 0.0000 | 0.99 |
| WC male | -0.0031 | 1.6x10<sup>-3</sup> | **-0.0128** | **5.4x10<sup>-9</sup>** | **0.0374** | **1.1x10<sup>-51</sup>** | **-0.0102** | **2.8x10<sup>-15</sup>** | 0.0007 | 0.80 |
| WHR | 0.0014 | 0.05 | -0.0016 | 0.30 | **0.0229** | **3.7x10<sup>-39</sup>** | **0.0051** | **1.6x10<sup>-8</sup>** | 0.0016 | 0.43 |
| WHR female | 0.0027 | 4.0x10<sup>-3</sup> | 0.0010 | 0.62 | **0.0221** | **2.8x10<sup>-22</sup>** | **0.0140** | **5.6x10<sup>-32</sup>** | 0.0027 | 0.31 |
| WHR male | 0.0003 | 0.74 | -0.0049 | 0.03 | **0.0242** | **1.4x10<sup>-21</sup>** | **-0.0059** | **7.6x10<sup>-6</sup>** | 0.0003 | 0.92 |

*Loci included in clusters:*
Beta Cell: *MTNR1B, CDKAL1, C2CD4A, HHEX, TCF7L2, SLC30A8, CDKN2A\_B, CDC123.CAMK1D, HNF1A, AP3S2, ZHX3, UBE2E2, ACSL1, PRC1, GIPR HNF1B, KCNJ11, KCNQ1\_2, ABO, ANK1, GLIS3, GLP2R, CTRB2, CDKN2A\_2 DUSP8, ADCY5, GIP, HNF4A, HSD17B12 TLE4*
Proinsulin: *ARAP1, SPRY2, DGKB\_2, IGF2BP2, CCND2, HNF4A, CDC123.CAMK1D*
Obesity: *FTO, MC4R, NRXN3, HSD17B12, RBMS1*
Lipodystrophy: *IRS1, GRB14, PPARG, LYPLAL1, ANKRD55, CMIP, KLF14, LPL, ANKRD55\_2, ARL15, ADCY5, C17orf58, POU5F1, MACF1, ZBED3, KIF9, ADAMTS9, CCND2, FAF1, MPHOSPH9*
Liver: *GCKR, CILP2, HLA.DQA1, PNPLA3, TSPAN8.LGR5*
*Abbreviations in text*
*Pvalues* < 2x10<sup>-4</sup> are **bolded**, representing a Bonferonni correction of 47 traits x 5 clusters.

**Fig. 1.**
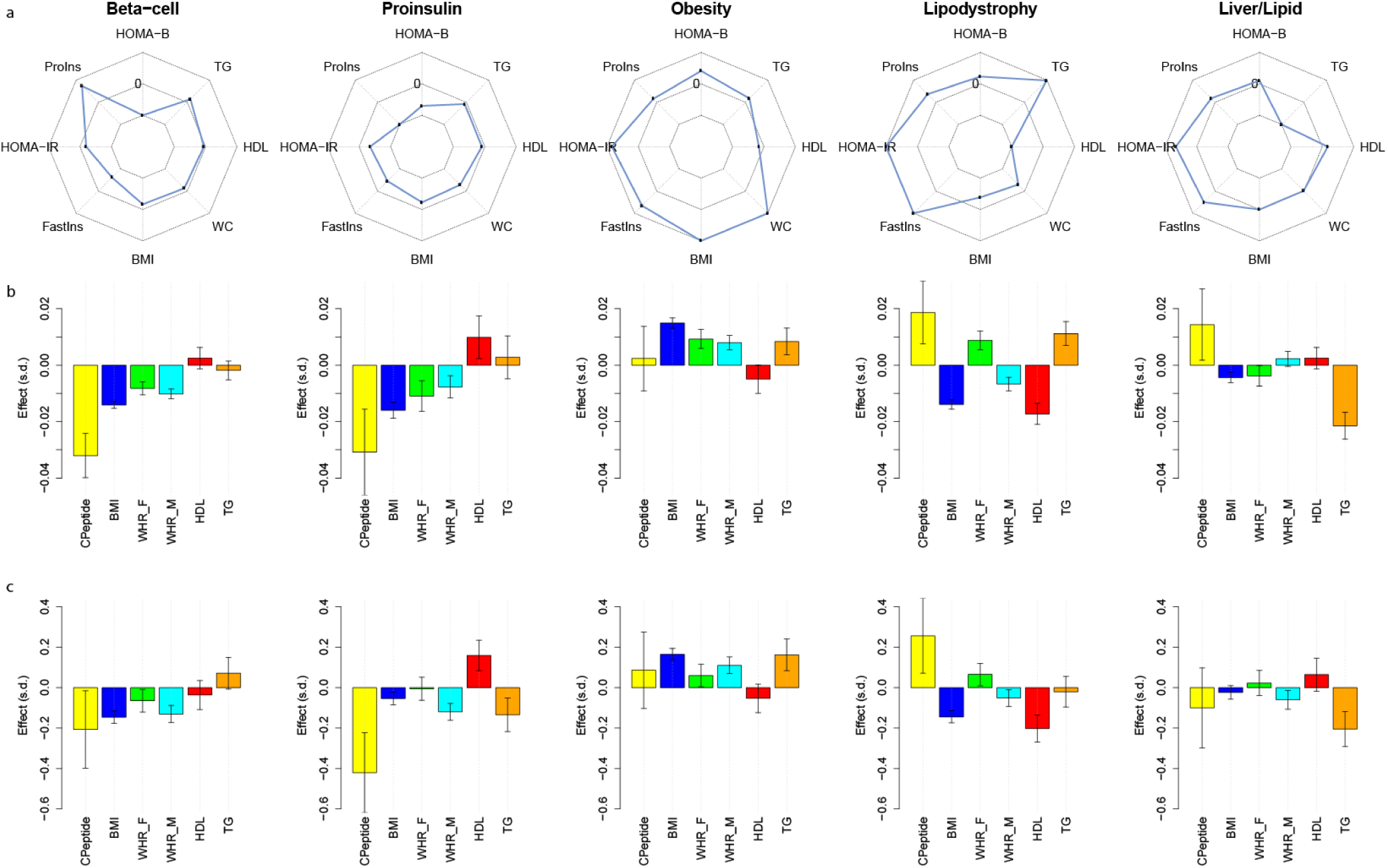
Cluster-defining characteristics. a. Association z-scores of cluster GRS’s and phenotypes derived from GWAS summary statistics shown in spider plot. The middle of the three concentric octagons is labeled “0,” representing no association between the cluster GRS and trait. Points falling outside the middle octagon represent positive cluster-trait associations, whereas those inside it represent negative cluster-trait associations. b. Associations of GRS’s in individuals with T2D with various traits. Results are from four studies (METSIM, Ashkenazi, Partners Biobank, and UK Biobank) meta-analyzed together. Effect sizes are scaled by the raw trait standard deviation. c. Traits of individuals with T2D who have GRS’s in the highest decile of a given cluster compared to all other individuals with T2D. Results are from the same four studies meta-analyzed together. Effect sizes are scaled by the raw trait standard deviation. Abbreviations: Proins – fasting proinsulin adjusted for fasting insulin, HOMA-B – homeostasis model assessment of beta cell function, TG – serum triglycerides, WC – waist circumference, BMI – body-mass index, Fastins – fasting insulin, HOMA-IR – homeostasis model assessment of insulin resistance, WHR-F – waist-hip-ratio in females, WHR-M, waist-hip ratios in males

The most strongly weighted traits for Clusters 1 and 2 relate to *insulin production and processing* in the pancreatic beta cell. In Cluster 1 (**Beta Cell cluster)**, these traits included decreased corrected insulin response (CIR), disposition index (DI), insulin at 30 minutes of OGTT (Ins30), beta-cell function by homeostasis model assessment (HOMA-B), and increased proinsulin levels adjusted for fasting insulin. Similarly, in Cluster 2 **(Proinsulin cluster)**, the top weighted traits included decreased Ins30 and HOMA-B, but also *decreased* proinsulin levels adjusted for fasting insulin, suggesting another mechanism impacting beta cell function. The loci that clustered most strongly into Cluster 1 include many well-known beta-cell-related loci: *MTNR1B*, *HHEX, TCF7L2, SLC30A8, HNF1A*, and *HNF1B* (Table 1, **Table S3**) (60, 61). Similarly, the two strongest weighted loci in Cluster 2 are *ARAP1*, a locus at which diabetes risk is thought to mediated by modulation of S*TARD10* expression in pancreatic beta cells (62), and *SPRY2*, a locus where the closest gene *SPRY2* regulates insulin transcription (63) **(Table S3)**. Using GWAS summary statistics, GRS’s composed of risk alleles (“GWAS GRS”) from top-weighted loci in each cluster (N loci Beta Cell=30, N loci Proinsulin=7; see Methods) were associated, as expected, with decreased HOMA-B (*P*-values <10^−10^) in both clusters and fasting insulin (*P*-values <5×10^−4^) in both clusters. The Beta Cell cluster GWAS GRS was associated with *increased* proinsulin (*P*<10^−9^), while the Proinsulin cluster GWAS GRS was associated with *decreased* proinsulin (*P*<10^−17^) (Table 1).

In contrast, Clusters 3, 4, and 5 all appeared to relate to mechanisms of *insulin response*. The traits that clustered most strongly with Cluster 3 (**Obesity cluster)** include increased waist circumference (WC), hip circumference (HC), BMI, and percent body fat. The top loci in this cluster include the well-known obesity-associated loci *FTO* and *MC4R* (Table 1, **Table S3**). The GWAS GRS for top-weight loci in this cluster (N loci=5) was significantly associated with these same traits (*P*-values <10^−24^) as well as increased fasting insulin unadjusted for BMI (*P*-value <10^−7^), but not fasting insulin adjusted for BMI (*P*-value =0.57). Thus, based on these association patterns, this cluster appeared to represent an obesity-mediated form of insulin resistance.

Cluster 4 (**Lipodystrophy cluster)** appears to represent the same “lipodystrophy-like” insulin resistance cluster previously suggested by Yaghootkar *et al*. (10, 64) with all the variants from that previous set which were also associated with T2D being among those loci most strongly weighted in this cluster (*PPARG*, *ANKRD55*, *ARL15*, *GRB14*, *IRS1*, and *LYPLAL1*) (Table 1, **Table S3**). This cluster’s strongest weighted traits were decreased insulin sensitivity index (ISI) adjusted for BMI, adiponectin, HDL, and increased triglycerides; GWAS GRS’s of alleles from the strongest weighted loci in this cluster (N loci=20) were associated with all of these traits (*P*-values <10^−20^). This cluster appears to represent a lipodystrophy or fat-distribution mediated form of insulin resistance. Interestingly, while an increased GWAS GRS from this cluster was significantly associated with increased waist-hip-ratio (WHR) in women (*P*<10^−31^), it was associated with decreased WHR in men (*P*<10^−5^). These ratios appear to be driven by GWAS GRS associations with decreased HC in both sexes (*P*-values <10^−9^), but a significant association with decreased WC only observed in men (*P*<10^−14^).

The final cluster (**Liver/Lipid cluster**) was notable for having decreased serum triglyceride levels, palmitoleic acid, urate, and linolenic acid, along with increased gamma-linolenic acid, as the traits most strongly weighted in this cluster. The GWAS GRS for the highest weighted loci (N loci=5) in this cluster were significantly associated with all the above traits (*P*-values <10^−7^). Notably, three of the top four weighted loci, *GCKR, TM6SF2*, and *PNPLA3* (Table 1, **Table S3**) have been previously associated with non-alcoholic fatty liver disease (NAFLD) (65), and functional work has implicated these loci in liver lipid metabolism (66–70).

#### Clusters are distinctly enriched for tissue enhancers or promoters

To gain further support for the suspected mechanistic pathways represented by these clusters and assess the biological distinctness of the clusters through an independent analysis, we assessed the top loci in each cluster for enrichment of epigenomic annotations (Figure 2). We expected each pathway to capture a different disease mechanism and thus localize largely to specific and distinct tissues; as common variants have been shown to lie typically in non-coding regions and presumably alter regulatory elements (enhancers, promoters), we assessed whether variants in the credible sets for loci in each cluster preferentially altered enhancers or promoters in specific cell types.

**Fig. 2.**
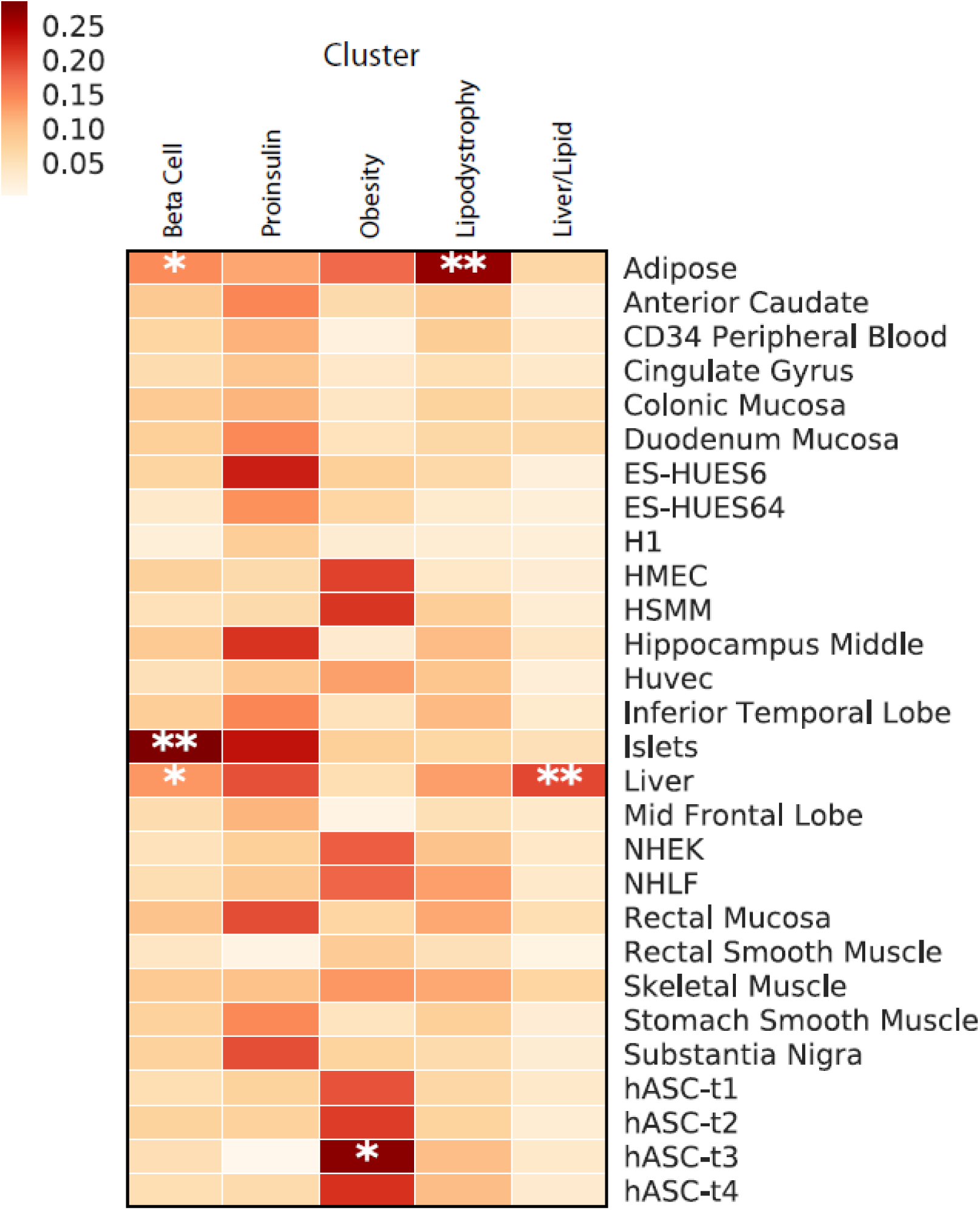
Enrichment for tissue-specific enhancers in clusters. Legend: Heatmap of associations of enrichment for enhancers and promoters from tissues residing within the top-weighted loci from each cluster. ** denotes P < 0.001, * P< 0.05. Cell line epigenetic data was obtained through the Epigenomics Roadmap; the authors played no role in the procurement of tissue or generation of the data for the two embryonic stem cell lines.

Within the Liver/Lipid cluster, there was significant enhancer/promoter enrichment in liver tissue (*P*<0.001), and within the Lipodystrophy cluster, there was significant enrichment in adipose tissue (*P*<0.001), each compared to the 27 other tissues assessed (Figure 2). The adipose tissue enrichment in the Lipodystrophy cluster was also the most significant across the five clusters (*P* <0.05) **(Figure S5)**. The loci in the Obesity cluster were most strongly enriched for enhancers/promoters in pre-adipose tissue, both compared to the other tissues and clusters (*P*-values <0.05). The two clusters suspected to be involved in beta-cell function (Beta Cell cluster and Proinsulin cluster), were most strongly enriched for pancreatic islet cell enhancers and promoters, with significant within (*P*<0.001) and across (*P*<0.05) enrichment for the Beta Cell cluster. Interestingly, the Proinsulin cluster had distinct enrichment in embryonic stem cells and brain tissue compared to the other clusters (*P*<0.05), which may reflect gene regulation related to early development, as beta cells and neurons have been thought to share gene expression patterns related to development (71). Thus in each case, the tissue enrichment supported the predicted biological mechanisms of each cluster.

#### Clusters are differentially associated with clinical outcomes from GWAS

To establish translational relevance, we next asked whether GWAS GRS’s from the strongest weighted loci in each cluster were associated with any clinical outcomes. We focused on clinical outcomes related to T2D available through GWAS: coronary artery disease (CAD), renal function as assessed by estimated glomerular filtration rate (eGFR) and urinary albumin:creatinine ratio (UACR), stroke risks, and blood pressure measures (Table 2). GWAS GRS’s from the Beta Cell and Lipodystrophy clusters were most strongly associated with increased risk of CAD (*P*-values <10^−7^). Increased Beta Cell cluster GWAS GRS was also significantly associated with increased risk for ischemic stroke (*P*=10^−4^), as well as stroke subtypes including large artery (*P*=10^−5^) and small vessel disease-related strokes (*P*=10^−4^), but not cardioembolic stroke (*P*=0.6). There were also nominally significant trends for these same stroke subtypes with the Lipodystrophy cluster. Only increased Lipodystrophy cluster GWAS GRS was significantly associated with increased blood pressure (SBP, beta = 0.07, *P*=6×10^−6^, DBP beta = 0.15, *P*=5×10^−9^, respectively). Renal function was most significantly associated with the Liver/Lipid cluster GWAS GRS; interestingly there was a significant association for increased Liver/Lipid GWAS GRS with reduced eGFR (beta = −0.002, *P*=10^−6^), but surprisingly also *reduced* UACR (beta=−0.009, *P*=10^−3^), whereas typically patients with diabetic kidney disease have reduced eGFR with *elevated* UACR. In contrast, increased Lipodystrophy cluster GWAS GRS was significantly associated with increased UACR (beta = 0.006, *P*=9×10^−5^). No cluster GWAS GRS was associated with risk of chronic kidney disease, defined as eGFR <60 ml/min/1.73 m^2^.

**Table 2.** Associations of cluster genetic risk scores and clinical outcomes from GWAS

|  | **Beta Cell**<br>N Loci = 30 |  | **Proinsulin**<br>N Loci = 7 |  | **Obesity**<br>N Loci = 5 |  | **Lipodystrophy**<br>N Loci = 20 |  | **Liver/Lipid**<br>N Loci = 5 |  | **Loci Combined**<br>N Loci = 62 |  |
| --- | --- | --- | --- | --- | --- | --- | --- | --- | --- | --- | --- | --- |
| **Outcome** | **beta** | **Pvalue** | **beta** | **Pvalue** | **beta** | **Pvalue** | **beta** | **Pvalue** | **beta** | **Pvalue** | **Beta** | **Pvalue** |
| CAD | **0.021** | **2.08E<sup>-12</sup>** | -0.003 | 0.67 | 0.016 | 0.04 | **0.021** | **2.5x10<sup>-8</sup>** | -0.009 | 0.27 | 0.017 | **1.2x10<sup>-15</sup>** |
| CKD | 0.003 | 0.35 | 0.009 | 0.22 | 0.015 | 0.04 | 0.0002 | 0.97 | 0.011 | 0.18 | 0.004 | 0.06 |
| eGFR | 0.000 | 0.87 | 0.0002 | 0.70 | -0.0008 | 0.06 | -0.00003 | 0.89 | **-0.002** | **1.2x10<sup>-6</sup>** | 0.000 | 0.07 |
| UACR | 0.001 | 0.42 | 0.003 | 0.27 | 0.003 | 0.32 | **0.006** | **9.0x10<sup>-5</sup>** | -0.010 | 3.7x10<sup>-3</sup> | 0.002 | 0.01 |
| Stroke_IS | **0.016** | **2.0x10<sup>-4</sup>** | 0.009 | 0.37 | 0.022 | 0.03 | 0.014 | 9.1x10<sup>-3</sup> | -0.002 | 0.84 | 0.013 | **1.3x10<sup>-5</sup>** |
| Stroke_CE | 0.004 | 0.59 | -0.001 | 0.94 | 0.048 | 0.01 | 0.005 | 0.62 | 0.002 | 0.93 | 0.007 | 0.20 |
| Stroke_LVD | **0.032** | **5.6x10<sup>-5</sup>** | -0.006 | 0.73 | 0.020 | 0.31 | 0.026 | 0.01 | -0.017 | 0.46 | 0.023 | **5.1x10<sup>-5</sup>** |
| Stroke_SVD | **0.032** | **2.6x10<sup>-4</sup>** | 0.029 | 0.17 | 0.027 | 0.22 | 0.036 | 1.5x10<sup>-3</sup> | -0.007 | 0.79 | 0.028 | **7.3x10<sup>-6</sup>** |
| SBP | 0.035 | 0.09 | -0.014 | 0.76 | -0.088 | 0.07 | **0.149** | **4.9x10<sup>-9</sup>** | -0.041 | 0.45 | 0.048 | 9.3x10<sup>-4</sup> |
| DBP | 0.014 | 0.30 | -0.027 | 0.36 | -0.046 | 0.14 | **0.073** | **6.4x10<sup>-6</sup>** | 0.0005 | 0.99 | 0.023 | 0.01 |

Abbreviations: coronary artery disease (CAD), chronic kidney disease (CKD), estimated glomerular filtration rate (eGFR), urinary albumin creatinine ratio (UACR), ischemic stroke all subtypes (IS), cerebroembolic (CE), large vessel disease (LVD), small vessel disease (SVD), systolic blood pressure (SBP), diastolic blood pressure (DBP)
Pvalues < 8x10<sup>-4</sup> are **bolded**, representing a Bonferonni correction of 10 outcomes x 6 groups.

#### Application of clusters to patients with T2D

To determine whether cluster GRS associations observed in large GWAS would have relevance to patients with T2D, we investigated cohorts of individuals with T2D ascertained from two epidemiological studies (METSIM [N=487] and the Diabetes Genes in Founder Populations (Ashkenazi) Study [N=509]) and two from biobanks (the Partners Biobank [N=2,065] and the UK Biobank [N=14,813]).

We first validated associations between cluster GRS’s and predicted phenotypes. In the up to 17,365 combined individuals with T2D from the four cohorts, increased individual-level GRS’s were associated with the expected salient traits (Figure 1b, **Table S6**): decreased BMI and percent body fat, (*P*-values <10^−20^) as well as fasting C-peptide (*P*=5×10^−5^) in the Beta Cell cluster; similarly decreased BMI (*P*=3×10^−8^) and fasting C-peptide (*P*=0.04) in the Proinsulin cluster; increased BMI, percent body fat, HC, and WC (*P*-values <10^−5^) in the Obesity cluster; decreased BMI, percent body fat, (*P*-values <10^−15^) and HDL (*P*=5×10^−5^) in the Lipodystrophy cluster; and decreased triglycerides in the Liver/Lipid cluster (*P*=2×10^−5^). Interestingly, as was noted with the GWAS GRS, a higher individual-level Lipodystrophy GRS had sex-differential associations with anthropometric traits: increased WHR in women (P=0.007), but decreased in men (*P*=0.005). A similar discrepant pattern was also seen with BMI adjustment **(Table S6)**.

We next asked whether this genetic approach could be utilized to identify individuals with T2D who had cluster-specific characteristics, in an initial attempt to stratify the population. In other words, would individuals with the largest GRS’s for each cluster differ from each other and all other individuals with T2D with regards to any clinical traits?

Consistently across the studies (N T2D=17,365), we observed ~30% of individuals with a GRS at the top 10^th^ percentile of just one cluster **(Table S8)**, which is what is expected by chance (under a binomial distribution). These individuals with the highest GRS’s differed significantly from all other participants with T2D (Figure 1c, **Table S7, Table S9)**: for instance, compared to all individuals across the three studies with T2D, those with extreme GRS in the Beta Cell cluster (N=1068) had decreased BMI, HC, and WC (*P-*values <10^−3^), and percent body fat (*P* <0.05) with a trend toward decreased fasting C-peptide (*P*=0.19), and those in the Proinsulin cluster (N=1,117) had significantly decreased fasting C-peptide levels (*P*=0.003); those in the Obesity cluster (N=1,206) had increased BMI, percent body fat, HC, and WC (*P-*values <0.05); those with extreme GRS in the Lipodystrophy cluster (N=1,134) had significantly decreased HDL, percent body fat, and BMI (*P*-values <0.01) and those with extreme GRS in the Liver/Lipid cluster (N=924) had significantly decreased triglycerides (*P*=0.01). Thus, individuals with T2D and a GRS uniquely at the top 10^th^ of one cluster as a group had representative trait characteristics distinguishing them from all other individuals with T2D (Figure 1c, **Table S7, Table S9**).

## Discussion

T2D, typically defined as hyperglycemia that is not autoimmune or monogenic in origin, is commonly recognized as a heterogeneous conglomerate of various pathogenic mechanisms, and therefore is unlikely to represent a single disease process. However, understanding of the biological pathways causing T2D to inform clinical management remains incomplete. Furthermore, despite over 100 T2D loci now identified, the relationship of these loci to disease pathways remains largely opaque.

Our work described here is the most comprehensive assessment of T2D loci clustering, including variant-trait associations for 94 T2D genetic loci and 47 diabetes-related metabolic traits in publically available GWAS datasets. We identify five robust clusters of T2D variants, which appear to represent biologically meaningful distinct pathways. The first two clusters (Beta Cell and Proinsulin) relate to pancreatic beta-cell function and differ most notably in the direction of association with proinsulin; both contain loci for which functional work has implicated beta-cell function in the causal mechanisms (e.g. (60, 61)). Additionally, the loci in both these clusters were highly enriched for positional overlap with pancreatic islet tissue enhancers and promoters. The three other clusters (Obesity, Lipodystrophy, Liver/Lipid) appear to represent different pathways causing insulin resistance: obesity-mediated, lipodystrophy (fat-distribution)-mediated, and liver-lipid-metabolism mediated. The Liver/Lipid cluster contains three of the top loci associated with NAFLD (65), and functional work has implicated these loci in liver lipid metabolism resulting in sequestration of lipid in the liver, resulting in decreased observed serum triglyceride levels (66–70). Additionally, these three clusters related to insulin action (Obesity, Lipodystrophy, Liver/Lipid) are enriched for variants overlaying enhancers in tissues that biologically support their proposed mechanisms: pre-adipocytes, adipocytes, and liver tissue respectively. GRS’s of top-weighted loci from the five clusters were also associated with particular clinical outcomes including increased systolic blood pressure, risk of coronary artery disease, and stroke, assessed using GWAS summary statistics.

Previous clustering efforts of T2D loci included less than half as many diabetes-related traits (6, 9, 11) and focused predominantly on unsupervised hierarchical clustering, a method that involves “hard clustering,” whereby a locus can be a member of only one cluster. Our analysis uses a novel clustering method bNMF, to enable a “soft-clustering”, whereby a variant can be a member of more than one cluster and also a more objective method for determining the number of clusters. With this method, the derived clusters are biologically interpretable and include mechanistic processes not previously captured by previous hard-clustering efforts, such as the Liver/Lipid cluster.

Our study additionally asks whether the clusters of variants have relevance to individuals with T2D. In four cohorts with up to 17,365 individuals with T2D of European ancestry, we show that individual level GRS’s for each cluster are associated significantly with predicted traits. Additionally, individuals with a very high GRS uniquely in the top 10^th^ percentile of one cluster had clinical features significantly distinguishing themselves from all other individuals with T2D; we observe consistently that this group comprises ~30% of persons with T2D, consistent with chance expectation, and importantly representing a sizable proportion of individuals with T2D.

Thus, these results suggest that genetics can be used to stratify a reasonable proportion of individuals with T2D who potentially belong to clinical subgroups. Such individuals could be classified based on their genetics and targeted for precision surveillance and therapeutics, should future studies find that these individuals differentially respond to medical interventions or confirm risk of particular clinical outcomes. Of course the threshold we chose of top 10^th^ percentile for each cluster GRS is arbitrary and further work is needed to determine clinically relevant thresholds or combinations of GRS from multiple clusters. Using the 10^th^ percentile cut-off, study participants with extreme GRS’s on aggregate had clinical characteristics distinguishable from others with T2D, however, at the individual-level, such clinical features might not be recognizable to a clinician due to the subtleties of the phenotypic characteristics, small differences in effects sizes, and/or potential for environmental influences. Furthermore, as germline genetic variation is constant throughout the lifetime and essentially unaltered by medications, it may provide a more robust metric than other biomarkers (which are contingent on environmental changes) on which to anchor an initial stratification step.

Other efforts have tried to identify subtypes of T2D patients (3, 4). In the most recent assessment of new-onset patients with T2D in Scandinavia, Ahlqvist and colleagues used phenotypic information to define five subgroups of diabetes: an autoimmune form (capturing type 1 diabetes and latent autoimmune diabetes in the adult), two severe forms (severe insulin deficient and severe insulin resistant diabetes), and two mild forms (obesity and age-related diabetes) (4). Importantly, in contrast to our clusters of genetic loci, these clusters are defined using clinical data and biomarkers at the time of diabetes diagnosis, and thus analysis of the same variables at a different time in the disease course or after treatment could yield inappropriate results. The T2D subtypes described by Ahlqvist *et al* seemed to have different genetic architectures (4), so we were interested to determine whether our clusters of genetic loci corresponded. Of the ten variants the authors found to be associated at nominal significance with their severe insulin deficient cluster (SIDD) and also included in our analysis, seven of these variants (or a proxy) had their strongest weights in our Proinsulin or Beta Cell clusters. Our Obesity cluster may correspond to the mild obesity-related diabetes of Ahlqvist *et al*., however none of the top-weighted variants in this cluster were included in their analysis. To the extent that severity of disease might be correlated with duration of exposure (with genetic exposure present at conception), our insulin resistance-related clusters might correspond to the severe insulin resistant diabetes (SIRD) of Ahlqvist *et al;* there were four variants found by the authors to be associated with the SIRD cluster at nominal significance, all of which were included in our analysis, and had their strongest weights in clusters we believe to relate to insulin resistance (Liver/Lipid and Lipodystrophy clusters). Interestingly, variants from several of our clusters were associated with the age-related diabetes cluster from Ahlqvist *et al*.

Beyond clarifying disease causal mechanisms and offering the potential for patient stratification, identification of the biologically-motivated clusters of T2D loci in our study may help also implicate loci with unknown function into pathways. For example, it is interesting that the *HLA.DQA1* locus is most highly weighted in the Liver/Lipid cluster; one might expect it to have a predominant autoimmune mechanism of action given its chromosomal location in the HLA region. The function of this locus remains to be discovered, however, our results suggest that it is more likely to influence insulin resistance than insulin deficiency. Membership to clusters may thus facilitate characterization of loci, which are generally challenging to study functionally.

Several loci included in this analysis have multiple independent signals reported in them (6, 8). We included 15 variants from six such loci (*ANKRD55 – two variants, DGKB – two variants, CDKN2A – three variants, KCNQ1 – four variants, CCND2 – two variants*, and *HNF4A – two variants*), offering an opportunity to see whether distinct signals from the same locus would cluster together **(Table S1)**. The multiple variants in *ANKRD55, CDKN2A, KCNQ1*, and *CCND2* mapped most strongly to the same cluster for each locus. At the *DGKB* locus, the signal represented by rs10276674 was most strongly weighted in the Proinsulin cluster, whereas the signal represented by rs2191349 was strongly weighted in the Beta Cell and Liver/Lipids clusters. In *HNF4A*, rs4812829 was most strongly weighted in the Beta Cell and Proinsulin clusters, whereas rs1800961 was most strongly weighted in the Lipodystrophy, Proinsulin and Liver/Lipid clusters **(Table S3)**. Interestingly, therefore, for *HNF4A*, the two signals separate into predominant insulin-deficiency and insulin resistance-related mechanisms. Potentially, therefore, cross-phenotype analysis can provide additional support for existence of independent signals at these loci, perhaps indicating tissue-specific regulatory mechanisms.

The strengths of this study include the novel application of a Bayesian form of NMF clustering to complex disease genetics. The clusters resulting from bNMF are more readily interpretable than hierarchical clustering, given that bNMF also clusters traits. Furthermore, allowance of an element to be part of more than one cluster (soft clustering) fits with our biological understanding of disease-causing genetic variants, whereby a particular variant may impact more than one biological pathway.

Limitations of this study include clustering of only available phenotypes from available GWAS. It is possible that future inclusion of additional traits would impact the clustering results, potentially even creating a new cluster for a mechanistic pathway not currently captured with available phenotypes. Additionally, we have focused on variants associated with T2D and related traits in populations of European ancestry. With additional studies from populations of diverse ethnic backgrounds, it would be ideal to include additional T2D-associated variants that were not included in the current analysis. Furthermore, the impact of the clusters on outcomes such as stroke and CAD was assessed through GWAS GRS using GWAS summary statistics, which came from studies including individuals with and without diabetes. It will be important to assess the association of cluster GRS and these outcomes in future work using individual level data; such analyses would ideally involve cohorts with large sample sizes and well-phenotyped outcomes. While NMF provides a very attractive method for variant-trait clustering, it is currently uncertain whether all weights or a thresholded approach is ideal for assignment of elements in a cluster. For our analysis, we developed a framework to do determine a reasonable threshold, however, this question could benefit from additional research.

In summary, clustering of genetic variants associated with T2D has identified five robust clusters with distinct trait associations, which likely represent mechanistic pathways causing T2D. These clusters have distinct tissue specificity, and patients enriched for alleles in each cluster exhibit distinct predicted phenotypic features. We observe a substantial fraction (~30%) of individuals with T2D who have T2D genetic risk factors highly loaded (in top 10^th^ percentile) from just one of the five clusters. It will be exciting to explore whether such individuals respond differentially to medications based on the pathway predominantly disrupted or have a differential rate of disease progression and diabetic complications. Classification of patients using data from designated genetic pathways may offer a step toward genetically informed patient management of T2D and improved individualized of care.

## Supplementary Materials

Fig. S1: Loci Associations to Clusters

Fig. S2: Trait Associations to Clusters

Fig. S3: Distribution of cluster weights for variants

Fig. S4: Deltas of cluster weights

Fig. S5: Enrichment for tissue-specific enhancers and promoters compared across clusters

Table S1: List of SNPs

Table S2: GWAS datasets

Table S3: Clustering results for T2D Loci

Table S4: Clustering results for traits

Table S5: GWAS GRS associations with traits

Table S6: Meta-analysis of GRS associations with traits in 4 studies of patients with T2D (adjusted for age and sex) plus sex-specific analyses

Table S7: Meta-analysis of comparison of mean trait in individuals with top 10% decile of one cluster GRS (cluster extreme) vs all subjects with T2D in 4 studies

Table S8: Counts of individuals from 5 studies in uniquely one top 10% decile of cluster GRS’s (cluster extreme)

Table S9: Comparison of median values of traits each top 10% decile of cluster GRS’s (cluster extremes) and in all others with T2D in two biobanks

## Acknowledgements

We wish to thank the thousands of volunteers who participated in the studies included in this work. Additionally we acknowledge the T2D-GENES initiative.

## Author contributions

MSU and JCF contributed with the conception of the work. MSU, CDA, MB, ML, GA, BG, and JCF contributed with data collection. MSU, JK, MvG, SBG, JMM, JC, JC, KG, JF, and GG contributed to data analysis. MSU, KG, JF, and JCF drafted the article. All authors contributed in the interpretation of data and revision of the article. All authors gave final approval of the version to be published.

